# Amino acid sensor conserved from bacteria to humans

**DOI:** 10.1101/2021.05.05.442820

**Authors:** Vadim M. Gumerov, Ekaterina P. Andrianova, Miguel A. Matilla, Annette C. Dolphin, Tino Krell, Igor B. Zhulin

## Abstract

Amino acids are recognized as signals by various receptors in bacteria, archaea, and eukaryotes. However, no common mechanism for amino acid recognition is currently known. Here we show that a subclass of a ubiquitous extracellular domain dCache_1 contains a simple amino acid recognition motif, and it is found throughout the Tree of Life. In bacteria, this motif exclusively binds amino acids, including GABA, and it is present in all major receptor types. In humans, this motif is found in α2δ subunits of voltage-gated calcium channels that are implicated in neuropathic pain and neurodevelopmental disorders. Our findings suggest that GABA-derived drugs bind to the same motif in human α2δ subunits that binds natural GABA ligands in bacterial chemoreceptors.

## MAIN TEXT

Amino acids are the building blocks of life and are involved in a variety of cellular processes including signal transduction. They serve as signals for various pathways in both prokaryotes and eukaryotes (*1*). Extracellular amino acids and their derivatives are recognized by dedicated receptors, such as G-protein coupled receptors (GPCRs) and ligand-gated ion channels in eukaryotes (*2, 3*) and chemoreceptors in bacteria and archaea (*4, 5*). In eukaryotes, Class C GPCRs, including GABA and metabotropic glutamate receptors, bind amino acid ligands at their Venus flytrap domain (*6*), whereas in ligand-gated ion channels, such as glycine and GABA receptors, amino acid ligands bind to an unrelated beta-sandwich-like domain (*7*). In bacterial chemoreceptors, amino acids are also recognized by unrelated ligand-binding domains, e.g. four-helix bundle (*8*) and dCache_1 (*9*). No common mechanism of amino acid sensing that would be present in all domains of life is currently known. Here we identify a simple conserved motif in the dCache_1 domain, which provides a common molecular mechanism for amino acid sensing for different types of receptors across the Tree of Life. The dCache_1 domain is the largest family of the Cache superfamily - ubiquitous extracellular ligand-binding sensors in bacteria and archaea that are also found in eukaryotes (*10, 11*). dCache_1 domains serve as sensory modules in all major types of bacterial and archaeal signal transduction systems (e.g. chemoreceptors, histidine kinases, diguanylate cyclases and phosphodiesterases, serine/threonine kinases and phosphatases) and they are also present in eukaryotic voltage-gated calcium channel (VGCC) α2δ subunits. In bacteria, dCache_1 domains bind various ligands, including amino acids, sugars, organic acids, and nucleotides (*5*), but their function in archaea and eukaryotes is unknown.

In a previous study, we showed that amino acid residues that are involved in binding amino acid ligands by dCache_1 domains from PctABC chemoreceptors in a bacterium *Pseudomonas aeruginosa* are conserved in many homologous chemoreceptors from gammaproteobacteria (*9*). In the present study we found that the same positions are conserved in all dCache_1 domains that are known to bind amino acids, whereas this conservation is lost in dCache_1 domains that bind ligands other than amino acids (Fig. 1A-B). Based on sequence and structure analysis, we propose the consensus amino acid binding motif (AA_motif) in dCache_1 domains (Fig. 1C, supplementary text, section 1), where Y121, R126 and W128 (from here and throughout the text, all motif positions are numbered according to *P. aeruginosa* chemoreceptor PctA, accession number NP_252999.1) make key contacts with the carboxyl group of the ligand and Y144 and D173 make key contacts with its amino group (Fig. 1D), as demonstrated for chemoreceptors from *P. aeruginosa* (*9*), *Campylobacter jejuni* (*12*), and *Vibrio cholerae* (*13*). R126, W128, and the Y129, were also proposed as conserved determinants of amino acid binding by others (*12*). To further verify the role of the AA_motif in amino acid binding, we mutated the key residues in the dCache_1 domain of the *P. aeruginosa* chemoreceptor PctA. The R126A substitution led to 61-fold decrease in the ligand binding affinity by PctA, whereas the D173A substitution completely abolished ligand binding (Fig. 1E, fig. S1). Similarly, mutations in these positions in the Tlp3 chemoreceptor in *C. jejuni* and in the Mlp37 chemoreceptor in *V. cholerae* significantly diminished amino acid binding (*12, 13*). Mutations in other positions of this motif also had a strong negative effect on amino acid binding in other bacterial receptors (table S2). Consequently, we renamed the AA_motif-containing dCache_1 domains to dCache_1AA.

**Fig. 1.**
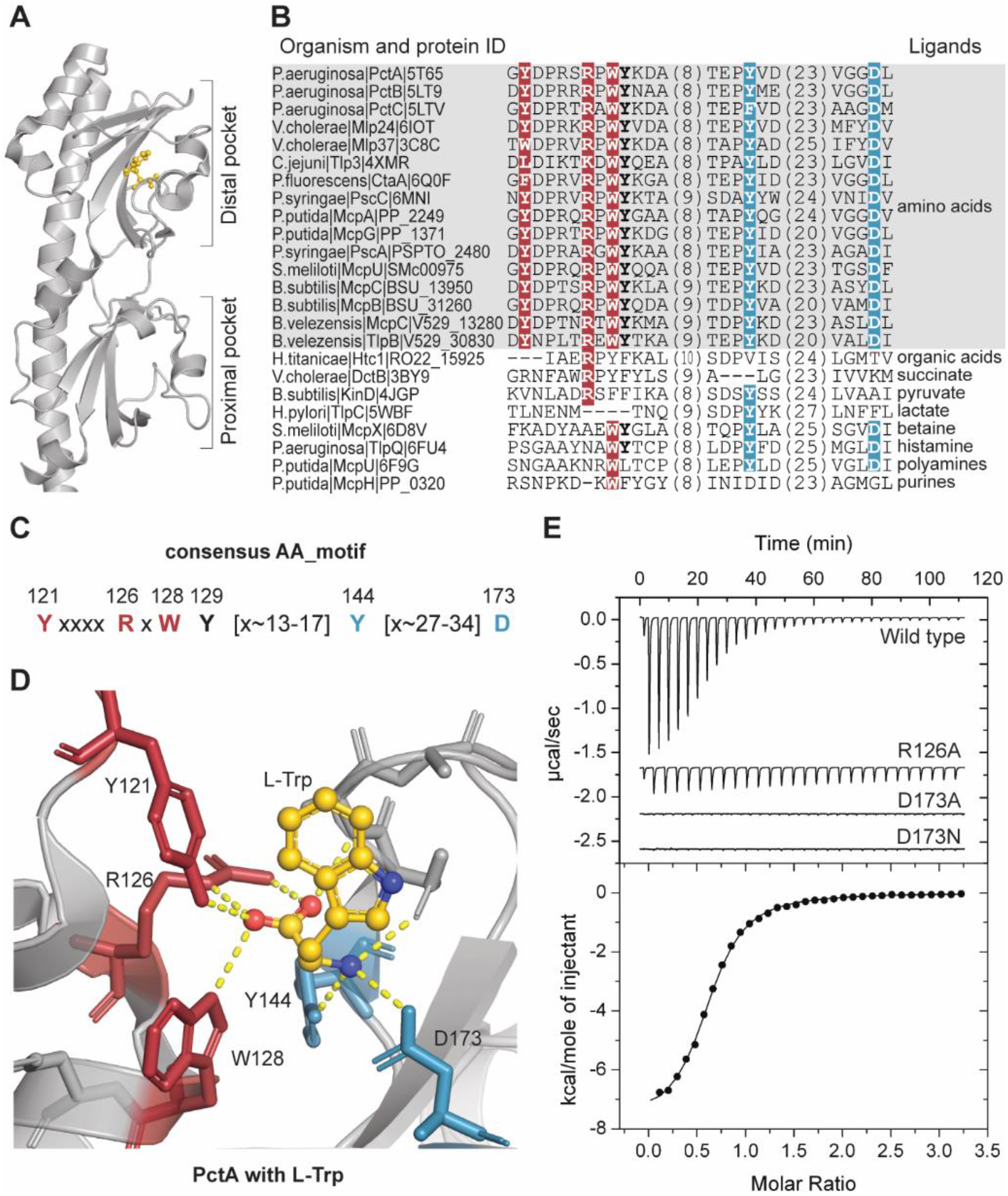
Amino acid binding motif. (**A**) dCache_1 domain of PctA chemoreceptor from *P. aeruginosa* PAO1 with bound L-Trp (gold) (PDB ID: 5T7M). (**B**) Protein sequence alignment of experimentally studied bacterial dCache_1 domains with respective ligands. AA_motif (in bold) is present in all amino acid binding dCache_1 domains (grey background). (**C**) Consensus amino acid binding motif. Numbers above the motif correspond to positions in PctA. (**D**) L-Trp interactions with AA_motif residues in the ligand binding pocket of PctA. (**B** to **D**) Here and in all figures: red, residues that coordinate the carboxyl group of the ligand; blue, residues that make contacts with the amino group. (**E**) Isothermal titration calorimetry study of L-Ala binding to wild type and mutated dCache_1 domain of PctA.

The list of dCache_1AA domains that are known to bind amino acids (Fig. 1B and table S2) contains no archaea or eukaryotes and has representatives of only three bacterial phyla (Proteobacteria, Campylobacterota, and Firmicutes) out of 111 phyla defined by the latest bacterial taxonomy (*14*). Consequently, we searched for the presence of dCache_1AA domains throughout the Tree of Life. First, we used the generalized AA_motif definition to search a dataset of 31,910 representative bacterial and archaeal genomes from the Genome Taxonomy Database (release 95) (*14*) (see Materials & Methods). This search identified the AA_motif in most bacterial and archaeal phyla for which at least ten high-quality genomes (>90% completeness) were available, and in all instances, this motif was located within the dCache_1 domain (table S4). Certain variability within the AA_motif permitting amino acid binding (table S2) was observed primarily in paralogous sequences (supplementary text, section 1). In addition, we found that ∼6% of motif containing sequences have a D173N substitution. To verify whether such substitution leads to the lack of amino acid binding we introduced it in *P. aeruginosa* PctA and showed that this change leads to a loss of function (Fig. 1E, supplementary text, section 2). As a result, we identified dCache_1AA domains in ∼11,000 proteins from the majority of bacterial and archaeal phyla (table S4), including important human pathogens, such as *Yersinia pestis, Vibrio cholerae, Clostridium botulinum, Legionella pneumophila*, and *Treponema pallidum*. dCache_1AA domains were found not only in chemoreceptors, but also in sensor histidine kinases, diguanylate cyclases and phosphodiesterases, serine/threonine kinases and phosphohydrolases and other proteins involved in signal transduction (table S5).

In the next step, we searched for amino acid sensing dCache_1 domains in eukaryotes. Cache domains were initially identified only in metazoan VGCC α2δ subunits (*10*), where they were described as “unusual”, “circular permutations”. Later, these were re-classified as dCache_1 domains with “uncertain boundaries” and detected in some other eukaryotic signal transduction proteins (*11*), however, no ligands were known to bind to these domains. In humans, α2δ Subunits are widely expressed in both the central and peripheral nervous systems and are implicated in various disorders, including schizophrenia, bipolar disorder, autism spectrum disorders, epilepsies, and neuropathic pain (*15, 16*). The α2δ-1 and α2δ-2 subunits bind GABA derived drugs gabapentin, pregabalin, and mirogabalin, of therapeutic benefit in neuropathic pain conditions (*17*). Coincidentally, GABA is a natural ligand for dCache_1AA domains of several bacterial chemoreceptors (table S2); however, it is unknown whether GABA derived drugs bind to the dCache_1 domain, and the precise location of this domain in α2δ is also unknown. In order to find out whether eukaryotic dCache_1 domains might contain the AA_motif, we performed a search initiated with this motif against several databases (see supplementary materials), which indeed retrieved several hundred eukaryotic sequences including α2δ subunits and the recently characterized CACHD1 proteins that also modulate VGCCs and are highly expressed in the thalamus, hippocampus, and cerebellum (*18, 19*).

In CACHD1, the AA_motif was mapped to a C-terminal region, where our analysis revealed a eukaryotic version of the dCache_1 domain (Fig. 2, supplementary text, sections 3, 7). No such motif was detected in the dCache_1 domain corresponding to VGCC_alpha2 in α2δ subunits (Fig. 2). Surprisingly, we found the first part of the AA_motif, YxxxxRxWY, in the domain currently annotated as VWA_N (a domain located N-terminally to the von Willebrand factor type A domain, Pfam accession number PF08399). Subsequent alignment of α2δ and CACHD1 with bacterial dCache_1_AA showed that bacterial sequences are well aligned with two regions of α2δ and CACHD1 proteins that are separated by the VWA domain (fig. S2). In the region located downstream of the VWA domain, we identified the second part of the AA_motif, Y[x∼27-34]D (Fig. 2, fig. S2). We took advantage of the recently published structure of the rabbit VGCC and its α2δ-1 subunit (*20*) to scrutinize the α2δ structure in light of these findings. Our structural analyses and topology tracking (see supplementary text, sections 4 and 7) together with the sequence analysis presented above revealed that α2δ and CACHD1 proteins are comprised of two dCache_1 domains, one inserted into another (Fig. 2, fig. S3, S5). Furthermore, the VWA domain is inserted into the first dCache_1 domain resulting in splitting the AA_motif in α2δ subunits into two parts. Excised and concatenated VGCC dCache_1 domains perfectly align with bacterial dCache_1 domains and find them in simple BLAST searches (table S8). Remarkably, the two parts of the AA_motif that are separated by the VWA domain come together spatially inside the binding pocket of the folded protein (Fig. 2D; fig. S4).

**Fig. 2.**
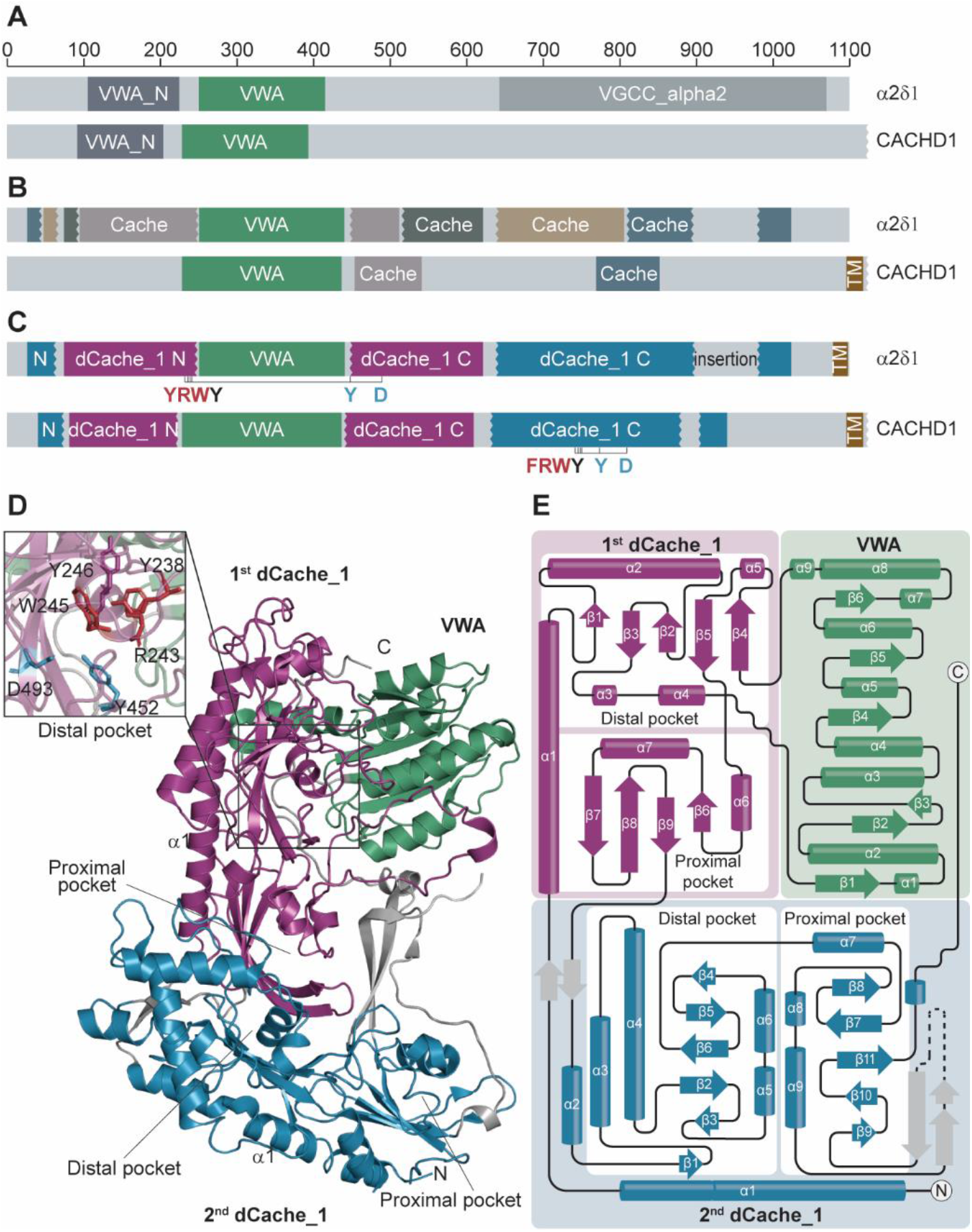
dCache_1AA domains in α2δ and CACHD1 subunits of VGCC. (**A-B**) Domains that are currently recognized in α2δ-1 and CACHD1 subunits by Pfam database (**A**) and experimental studies (*21*) and (*18*) (**B**). (**C**) Domain architectures of α2δ-1 and CACHD1 subunits revealed in the present study. (**D**) Structural composition of α2δ-1 subunit uncovered in the present study shown on the solved structure (PDB ID: 6JPA (*20*)). A close-up view of the dCache_1AA distal pocket (top left) showing the spatial proximity of the AA_motif residues despite the VWA insertion. (**D**) The α2δ-1 subunit topology shows that the VWA domain is inserted into the first dCache_1 domain, which in turn is inserted into the second dCache_1 domain.

R241A substitutions in the murine and porcine α2δ-1 (corresponds to R126 in PctA) were shown to completely abolish the ability to bind pregabalin and gabapentin, respectively (*22, 23*). Furthermore, the R241A substitution in the murine α2δ-1 has been shown to result in a significant decrease in divalent cation current through CaV2.2 channels (*23*). Leucine and isoleucine were shown to bind to α2δ-1 (*24*) and inhibit gabapentin binding by the subunit (*25*). To further explore whether the AA_motif in α2δ-1 might serve as a site for binding GABA derived drugs and amino acids we performed computational docking experiments with the available structure of the rabbit α2δ-1 protein (*20*). Results of computational experiments combined with structural superimpositions showed that α2δ-1 binds the drug molecules and amino acids by the AA_motif and the relative affinities agree with the available experimental data (Fig. 3 and supplementary text, sections 5-6). Position and orientation of the ligands in α2δ-1 and in PctA/PctC are very similar, and the ligands make polar contacts with the amino acid binding motif in a very similar fashion (Fig. 3, fig. S4), which is remarkable considering the evolutionary distance between bacteria and mammals.

**Fig. 3.**
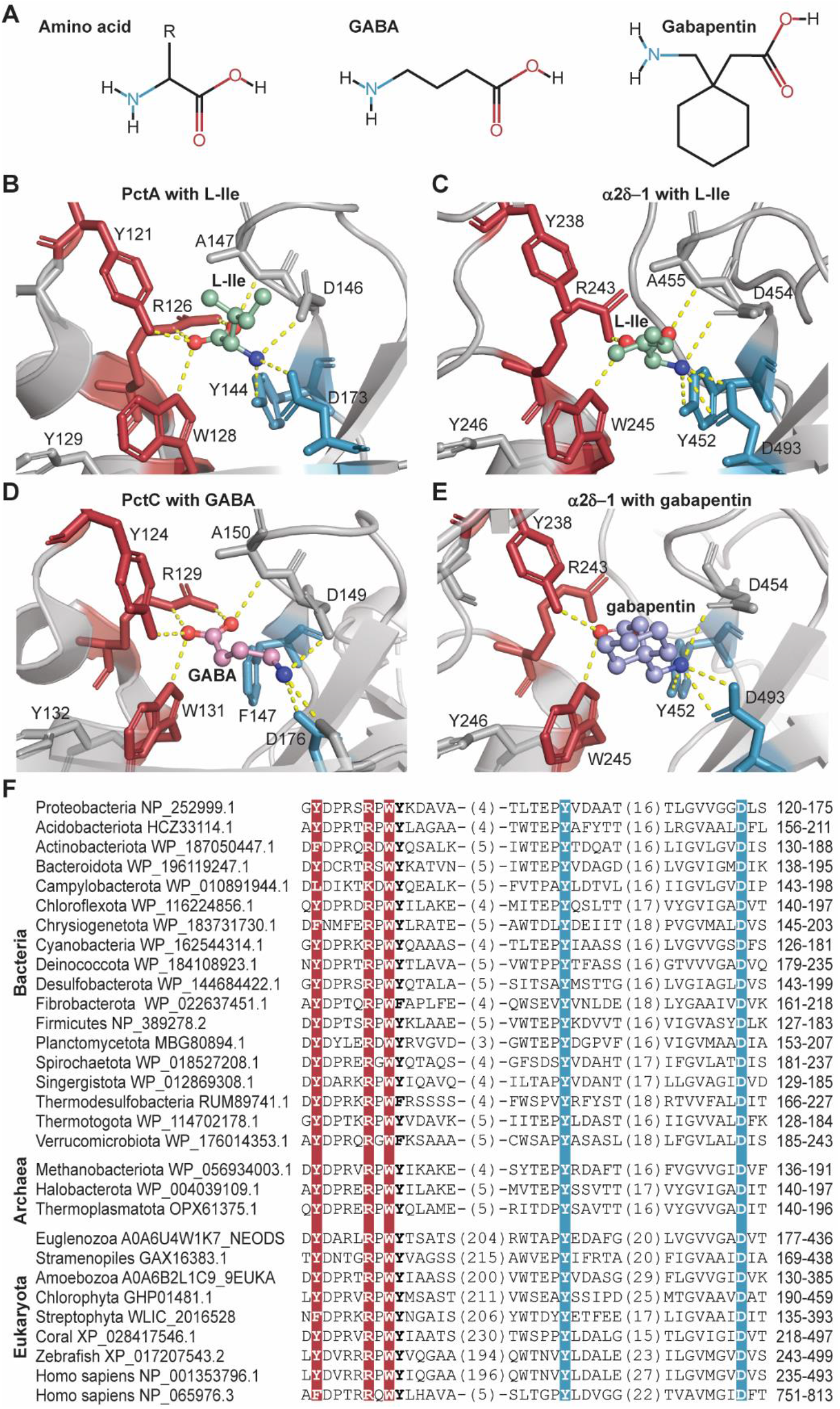
Both bacterial and mammalian receptors bind amino acid ligands through the conserved AA_motif. (**A**) Structural comparison of the ligands found to bind dCache_1AA. (**B-E**) Ligand binding modes of bacterial and eukaryotic dCache_1AA: PctA with L-Ile (**B**, PDB ID: 5T65), α2δ-1 with docked L-Ile (**C**), PctC in complex with GABA (**D**, PDB ID: 5LTV), and α2δ-1 with docked gabapentin (**E**). (**F**) Protein sequence alignment of the dCache_1AA from major phyla of Bacteria, Archaea and Eukaryota shows that AA_motif is conserved throughout the Tree of Life.

To establish the prevalence of the motif in Eukaryota and its evolutionary history we analyzed available eukaryotic genomes (supplementary text, section 8). We found that the dCache_1AA containing homologs of α2δ and CACHD1 proteins are universally present in eukaryotes except for flowering plants, fungi, and two protozoan lineages, in which the proteins were presumably lost (Fig. 4, supplementary text, section 8). A dCache_1AA domain containing protein was likely present in the Last Eukaryotic Common Ancestor (LECA) (fig. S7, table S7) conceivably as a result of horizontal gene transfer from bacteria. The VWA domain has been inserted into the first dCache_1 domain as early as LECA (see supplementary text, section 8), while the first dCache_1 domain has been inserted into the second domain in one eukaryotic branch prior to Choanoflagellata divergence. Following this insertion, the protein has undergone multiple duplications in Metazoa and has given rise to four α2δ paralogs and one CACHD1 protein in Vertebrata (fig. S6, S7). Our analysis also demonstrated that the first dCache_1 domain of α2δ subunits is under stronger selective pressure than the second one; in contrast, both dCache_1 domains of CACHD1 subunit are under strong selective pressure (supplementary text, section 9).

**Fig. 4.**
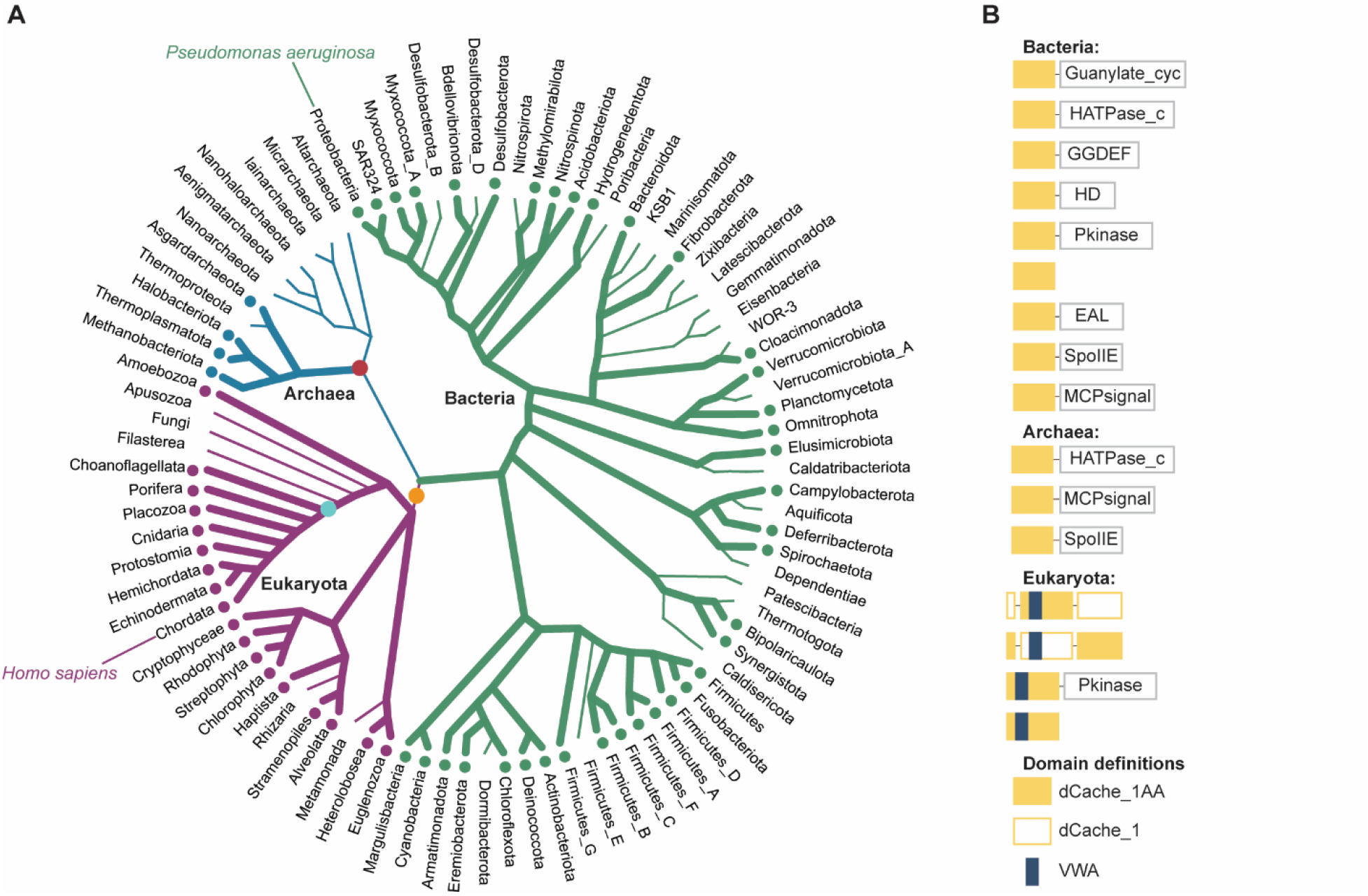
The AA_motif across the Tree of Life. (**A**) Distribution of the dCache_1AA across major lineages of life. Thick lines with dots at the tips denote the presence of the AA_motif. Red circle indicates horizontal gene transfer of the dCache_1AA to Archaea. Orange circle shows probable event of the horizontal transfer of dCache_1AA to Eukaryota and VWA domain insertion. Circle in teal indicates an event of insertion of 2^nd^ dCache_1 into the 1^st^ dCache_1 domain in eukaryotes. (**B**) Prevalent domain architectures of the dCache_1AA containing proteins found in each domain of life are shown (the Pfam domain nomenclature is used). See supplementary materials for details.

In this work we have described a universal amino acid binding sensor, which is present throughout the Tree of Life. Consequently, we assign specific biological function – amino acid sensing – to thousands of receptors in bacteria and archaea. It is especially important for human pathogens because amino acids are key mediators of pathogenicity (*26*). We identified the amino acid binding motif in α2δ and CACHD1 subunits of voltage-gated calcium channels and implicated it as the binding site for GABA-derived drugs in human α2δ subunits. This finding provides new opportunities for improving drugs targeting various neurobiological disorders.

