## Supplemental text, figures, and tables for "Amino acid sensor conserved from bacteria to humans"

**This PDF file includes:**

Materials and Methods
Supplementary Text
Figs. S1 to S7
Tables S1-3, S2, and S9

**Other Supplementary Materials for this manuscript include the following:**

Excel Tables S4 to S8
File 1

**Supplementary Text** sections:

1. Identification of the AA_motif in Prokaryota
2. Ligand binding studies
3. Identification of the AA_motif in Eukaryota
4. α2δ-1 subunit domain composition
5. The ligand-binding pocket of α2δ-1 contains the universal amino acid binding motif
6. α2δ-1 subunit and bacterial chemoreceptors bind ligands in a similar fashion
7. CACHD1 has the same domain architecture as α2δ-1 subunit
8. Evolution of amino acid binding dCache_1 domain in Eukaryota
9. N-terminal dCache_1 domain of α2δ subunits is under stronger selective pressure than the C-terminal dCache_1 domain

**Materials and Methods**

Identification and collection of prokaryotic amino acid binding motif-containing proteins.

Protein sequences of representative bacteria and archaea were downloaded from the Genome Taxonomy database (GTDB, Release 95.0) (*1*). Retrieved dataset was scanned with the dCache_1 Hidden Markov Model (HMM) profile obtained from the Pfam database (Release 34.0) (*2*) with the E-value threshold of 0.01 both for sequences and domains. Next, protein sequence regions corresponding to dCache_1 domain were extracted from the identified sequences.

To avoid the identification of false positive amino acid binding proteins we varied positions in the motif not within amino acid physicochemical groups but within those amino acids that were reported to permit amino acids binding (**Table S2**). The following AA_motif definition was used (in the regular expression notation): [YFWL]….[RK].W[WYF]{n1}[YF]{n2}[D]. In this expression, any amino acid inside the given brackets can be present in the corresponding position, n1 – a varying distance of approximately 13-17 amino acid residues in prokaryotic dCache_1 domains, n2 – a varying distance of approximately 27-34 amino acid residues in prokaryotic dCache_1 domains. In eukaryotic dCache_1 domains that have the von Willebrand factor type A (VWA) domain insertion, n1 is a varying distance of approximately 215-245 amino acid residues.

Next, we scanned the dCache_1 domain protein sequences encoded in the genomes of the representative set of bacteria and archaea with the first part of the motif, [YFWL]….[RK].[W][WYF], and extracted corresponding sequences. Using these sequences, we have built two multiple sequences alignments (MSAs): one – with the bacterial sequences and one – with the archaeal. Exploring the alignments, we identified positions corresponding to the amino acid binding motif. In the bacterial alignment coordinates corresponding to the motif are the following: Y(**1160**)….R(**1278**).W(**1290**)Y(**1291**){n1}Y(**1431**){n2}D(**1647**). In the archaeal alignment the coordinates are the following: Y(**163**)….R(**168**).W(**170**)Y(**171**){n1}Y(**189**){n2}D(**219**). Using the MSAs and the motif definitions and coordinates we calculated how many sequences have the second part of the motif ({n1}[YF]{n2}[D]): 10700 bacterial and 108 archaeal sequences. Next, using these sequences that have the motif in the full form we calculated the motif variant frequencies and abundances. Full protein sequence alignments can be found in https://github.com/ToshkaDev/Motif repository, “GTDB_Representative_Set” folder. Following this step, we retrieved full taxonomy information from the GTDB metadata tables (the tables are located here https://data.ace.uq.edu.au/public/gtdb/data/releases/release95/95.0/) for the genome assemblies in which we found the motif.

Additionally, the NCBI RefSeq (Release 202) and Uniprot (Release 2021_02) databases were scanned for the AA_motif containing proteins. Full alignments of dCache_1 domains with the AA_motif from these datasets are in https://github.com/ToshkaDev/Motif repository. We identified 32395 dCache_1 domain containing protein sequences with the AA_motif in RefSeq database and 11330 sequences in Uniprot.

The AA_motif positions in the RefSeq MSA:

Y(**2091**)....R(**2302**).W(**2385**)Y(**2414**){n1}Y(**2727**){n2}D(**3326**)

The AA_motif positions in the Uniprot MSA:

Y(**308**)....R(**313**).W(**315**)Y(**316**){n1}Y(**345**){n2}D(**420**)

Identification and collection of eukaryotic amino-acid binding proteins.

To identify dCache_1 domain containing proteins with the AA_motif in eukaryotes, human α2δ-1 and CACHD1 protein sequences (protein IDs are NP_001353796.1 and NP_065976.3, respectively) were used as queries for BLASTP and PSI-BLAST searches against NCBI RefSeq, NCBI Nonredundant, Uniprot (*3*), and 1KP (*4*) databases. The presence of the motif was verified by constructing multiple sequence alignments with already identified motif-containing eukaryotic and bacterial sequences. Eukaryotic protein multiple sequence alignment can be found in https://github.com/ToshkaDev/Motif repository. Taxonomy information was retrieved from the NCBI Taxonomy database (https://www.ncbi.nlm.nih.gov/taxonomy).

Construction of the Tree of Life

For the schematic representation of the Tree of Life (**Fig. 4**) bacterial and archaeal phylogeny was retrieved from the GTDB taxonomy. Phyla with at least 10 genomes were depicted. Eukaryotic phylogeny was adapted from (*5*) and (*6*). The overall tree topology is based on (*7*).

Multiple sequence alignment, phylogeny inference, and domains identification

MSAs were constructed using L-INS-i algorithm of MAFFT (*8*). Jalview (*9*) was used to explore and edit the alignments. Neighbor-joining and Maximum likelihood phylogenetic trees were built in TREND (*10*) applying JTT substitution model and 500 bootstrap resamplings.

Domains were identified running TREND with the Pfam HMM profiles. The generated data were downloaded in JSON format from the website and processed programmatically to determine domain architecture variants and abundances (**Table S5**). Additional sensitive profile-profile searches were carried out using HHpred (*11*).

Taxonomic trees were retrieved from Annotree (*12*) and the NCBI Taxonomy database and were edited in iTOL (*7*).

Computational Docking experiments

AutoDock Vina (*13*) was used for computational docking experiments. The α2δ-1 subunit structure was obtained from the rabbit voltage gated calcium channel structure (PDB ID: 6JPA, (*14*)). The subunit was prepared using MGLTools (*15*). For the experiments we downloaded ligands from the Zink database (*16*) in mol2 format and prepared them for the analysis using Open Babel toolbox (*17*) and custom shell script. The following settings were used to run the docking simulations:

a) coordinates of the center of the simulation box (Angstroms):

X: 198.681

Y: 165.755

Z: 221.688

b) the box dimensions (Angstroms):

X: 22

Y: 22

Z: 24

c) the search exhaustiveness: 8

Protein structures manipulations

The human α2δ-1 and CACHD1 subunits were modeled based on the solved rabbit α2δ-1 structure using I-TASSER (*18*) with default parameters. Protein structures were explored in PyMoL (*19*) and Python Molecular Viewer from the MGLTools package (*15*).

Site-directed mutagenesis

An overlapping PCR mutagenesis approach was employed to construct the substitution mutants of PctA-LBD using primers listed in **Table S1**. The resulting PCR fragments were cloned into the NdeI/BamHI sites of the expression vector pET28b(+). *Escherichia coli* DH5α was used as host for gene cloning. The resulting plasmids (**Table S1**) were verified by DNA sequencing. Mutant proteins were purified following the protocol for the wild-type protein.

Protein expression, purification, and isothermal titration calorimetry (ITC) analyses.

Native and mutant PctA-LBD proteins were overexpressed in *E. coli* BL21 (DE3) and purified as described in (*20*). Microcalorimetric analyses were conducted according to (*20*).

**Table S1. Strains, plasmids and oligonucleotides used in this study.**

| | **Relevant characteristics** | **Reference or source** |
|---|---|---|
| **Strains** | | |
| *E. coli* BL21 (DE3) | F$^-$, *ompI*, *hsdS*$_B$ (r$^-_B$ m$^-_B$) *gal*, *dam*, *met* | (*21*) |
| *E. coli* DH5α | *supE44 lacU169* (*Ø80lacZ*ΔM15) *hsdR17* ($r_k^- m_k^-$), *recA1 endA1 gyrA96 thi-1 relA1* | (*22*) |
| Plasmids | | |
| pET28b(+) | Km$^R$; Protein expression plasmid | Novagen |
| **pET28-PctA-**LBD | Km$^R$; pET28b(+) derivative containing DNA fragment encoding PctA-LBD | (*20*) |
| pMAMV359 | Km$^R$; pET28b(+) derivative containing DNA fragment encoding PctA-LBD (R126A) | Present study |
| pMAMV360 | Km$^R$; pET28b(+) derivative containing DNA fragment encoding PctA-LBD (D173A) | Present study |
| pMAMV376 | Km$^R$; pET28b(+) derivative containing DNA fragment encoding PctA-LBD (D173N) | Present study |

**Oligonucleotides**

| **Name** | **Sequence (5´-3´)** | **Purpose** | **Source** |
|---|---|---|---|
| PctA-LBD-F | GTTCACCCATATGAACGATTACCTGCAGCGCAACG | *pctA-LBD* into pET28b(+) | (*20*) |
| PctA-LBD-R | GCGGATCCTCAGGCCGAGACGCGGAACTTG | *pctA-LBD* into pET28b(+) | (*20*) |
| R126A-F | TCCGCGCAGCGCGCCCTGGTACAAG | Mutant R126A | Present study |
| R126A-R | CTTGTACCAGGGCGCGCTGCGCGGA | Mutant R126A | Present study |
| D173A-F | GTAGGCGGCGCCCTCAGCCTGAAG | Mutant D173A | Present study |
| D173A-R | CTTCAGGCTGAGGGCGCCGCCTAC | Mutant D173A | Present study |
| D173N-F | GTAGGCGGCAACCTCAGCCTGAAG | Mutant D173N | Present study |
| D173N-R | CTTCAGGCTGAGGTTGCCGCCTAC | Mutant D173N | Present study |

**Supplementary Text**

1. Identification of the AA_motif in Prokaryota

We collected information about experimentally studied dCache_1 domain containing proteins capable of amino acid binding (**Table S2**). Exploring retrieved protein sequences, we found that all of them contained the same conserved residues. Using available structures of dCache_1 domains in complex with amino acid ligands (**Table S2**) we established that a common pattern of amino acid binding exists. Remarkably, the same conserved residues of dCache_1 domains formed hydrogen bonds with either carboxyl or amino group of the amino acid ligand. Based on our previous study (*23*), protein sequence conservation, and structural analysis, we proposed the consensus amino acid binding motif (AA_motif) in the ligand binding pocket of dCache_1 domain: Y....R.WY[~13-17]Y[27-34]D (**Fig. 1, Table S2**). Further we noted that the AA_motif permits a certain variability within amino acid groups in each position (**Table S2**).

Using this data, we scanned GTDB and retrieved bacterial and archaeal protein sequence containing dCache_1 domain with the AA_motif (see **Materials and Methods**). We found dCache_1 domains with the AA_motif in 51 bacterial and 4 archaea phyla (**Table S4,** sheets 1 and 2) and counted the AA_motif variants (**Table S3;** Excel **Table S4,** sheets 3 and 4). Tracking the motif variability within genomes we established that the top three variants (Y....R.WY{n1}Y{n2}D, F....R.WY{n1}Y{n2}D, Y....R.WF{n1}Y{n2}D) are present almost universally in motif-containing genomes, while the most abundant consensus motif variant, Y....R.WY{n2}Y{n2}D, is present in 86% of the bacterial and 100% of the archaeal phyla (Excel **Table S4,** sheets 7 and 8). Other variants coexist along with these top variants, mostly in paralogous proteins (Excel **Table S4,** sheets 5 and 6).

**Table S2. Experimentally studied amino acid binding proteins containing dCache_1 domain with AA_motif.** For each protein its identifier, ligand repertoire, affinities, and amino acid binding motif variant is shown. For proteins with solved structure contacts with the ligand carboxyl and amino group are shown.

| Organism | Protein Name | Protein ID | Reference | PDB ID | Ligand | $K_D$ (μM) | **AA_motif;** hydrogen bonds with the ligand carboxyl group are in **red** and amino group are in **blue** | Mutagenic analysis |
|---|---|---|---|---|---|---|---|---|
| *Pseudomonas aeruginosa PAO1* | PctA | NP_252999.1 | (*20, 23-25*) | | L-Arg | 1.8 ± 0.4 | **Y....R.WY Y D** | Present study:<br><br>Mutation of Y....R.WY Y **D** to A abolished L-Ala and L-Ser binding<br><br>Mutation of Y....R.WY Y **D** to N strongly reduced L-Ala binding<br><br>Mutation of Y....**R**.WY Y D to A reduced L-Ala and L-Ser binding by 61- and 78-fold, respectively. |
| | | | | | L-Lys | 1.6 ± 0.1 | | |
| | | | | | L-Tyr | 5.2 ± 1.7 | | |
| | | | | 5T7M | L-Trp | 2.3 ± 0.9 | | |
| | | | | | L-Phe | 1.1 ± 0.2 | | |
| | | | | | L-Ala | 0.72 ± 0.1 | | |
| | | | | 5T65 | L-Ile | 24 ± 5.6 | | |
| | | | | | L-Leu | 116 ± 10 | | |
| | | | | 5LTX | L-Met | 0.91 ± 0.2 | | |
| | | | | | L-Asn | 2.0 ± 0.2 | | |
| | | | | | L-Ser | 1.2 ± 0.2 | | |
| | | | | | L-Cys | 0.79 ± 0.1 | | |
| | | | | | L-Thr | 0.28 ± 0.1 | | |
| | | | | | L-His | 28 ± 5 | | |
| | | | | | L-Pro | 0.60 ± 0.2 | | |
| | | | | | L-Gly | 21 ± 7.7 | | |
| | | | | | L-Val | 0.34 ± 0.2 | | |
| *P. aeruginosa PAO1* | PctB | NP_253000.1 | (*20, 24-26*) | 5LT9 | L-Arg | 64 ± 4 | **Y....R.WY Y D** | |
| | | | | | L-Lys | 1096 ± 88 | | |
| | | | | | L-Ala | 641 ± 50 | | |
| | | | | | L-Met | 46 ± 2 | | |
| | | | | 5LTO | L-Gln | 1.2 ± 0.1 | | |
| *P. aeruginosa* | PctC | NP_252997.1 | (*20,* | 5LTV | GABA | 1.2 ± 0.3 | **Y....R.WY F D** | |

| | | | | | | | | |
|---|---|---|---|---|---|---|---|---|
| *PAO1* | | | *24-26*) | | L-His | 17 ± 3 | | |
| | | | | | L-Pro | 80 ± 6 | | |
| *P. putida KT2440* | McpG | WP_010952482.1 | (*27*) | | GABA | 0.175 | **Y....R.WY Y D** | |
| *P. putida KT2440* | McpA | WP_010953225.1 | (*28*) | | Gly | 35 ± 2 | **Y....R.WY Y D** | |
| | | | | | L-Ala | 13 ± 0.6 | | |
| | | | | | L-Cys | 0.6 ± 0.1 | | |
| | | | | | L-Ser | 43 ± 2 | | |
| | | | | | L-Asn | 4.3 ± 0.2 | | |
| | | | | | L-Gln | 5.5 ± 0.2 | | |
| | | | | | L-Phe | 2.3 ± 0.1 | | |
| | | | | | L-Tyr | 12.1 ± 0.9 | | |
| | | | | | L-Val | 373 ± 42 | | |
| | | | | | L-Ile | 85 ± 5 | | |
| | | | | | L-Met | 5.8 ± 0.6 | | |
| | | | | | L-Arg | 1.2 ± 0.1 | | |
| *Sinorhizobium meliloti RU11/001* | McpU | WP_010968886.1 | (*29*) | | Pro | 104 | **Y....R.WY Y D** | mutation of Y....R.WY Y **D** to A abolished taxis toward Pro; mutation of Y....R.WY Y **D** to E showed intermediary response |
| *Sinorhizobium meliloti RU11/001* | McpU | WP_010968886.1 | (*30*) | | Arg | 350 | **Y....R.WY Y D** | |
| | | | | | Phe | 53 | | |
| | | | | | Trp | 34 | | |
| | | | | | Pro | 42 | | |
| *P. fluorescens Pf0-1* | CtaA | WP_011335661.1 | (*31*) | 6Q0F | L-Val | 4.7 ± 1.2 | **F....R.WY Y D** | |
| | | | | 6PXY | L-Ala | 5.2 ± 0.3 | | |
| | | | | 6PY5 | L-Ser | 9.5 ± 1.1 | | |
| | | | | 6PY4 | L-Leu | 11.9 ± 1.8 | | |
| | | | | 6Q0G | L-Pro | 13.5 ± 0.3 | | |
| | | | | 6PY3 | L-Ile | 27.4 ± 0.9 | | |
| | | | | | L-Arg | 446 ± 17.6 | | |
| *Bacillus velezensis SQR9* | McpC | AHZ15354.1 | (*32*) | | L-Leu | 3.6 ± 0.78 | **Y....R.WY Y D** | |
| | | | | | L-Pro | 3.6 ± 0.78 | | |
| *B. velezensis SQR9* | TlpB | AHZ17109.1 | (*32*) | | L-Phe | 3.17± 0.89 | **Y....R.WY Y D** | |

| | | | | | | | | |
|---|---|---|---|---|---|---|---|---|
| *B. subtilis OI1085* | McpB | WP_003243461.1 | (*33*) | | L-Asn | 14 | **Y....R.WY Y D** | mutation of **Y**....R.WY Y D to A showed defects in taxis toward Asp; mutation of **Y**....R.WY Y D to F had no effect |
| *B. subtilis OI1085* | McpC | WP_003245443.1 | (*34*) | | L-Pro | 14 | **Y....R.WY Y D** | |
| | | | | | L-Thr | 21 | | |
| | | | | | L-Ser | 90 | | |
| | | | | | L-Val | 154 | | |
| | | | | | L-Ala | 18 | | |
| | | | | | L-Tyr | 360 | | |
| | | | | | L-Phe | 492 | | |
| | | | | | L-Leu | 72 | | |
| | | | | | L-His | 320 | | |
| *Vibrio cholerae O395N1* | Mlp37 | AAF96820.1 | (*35*) | 5AVF | taurine | 3.2 | **W....R.WY Y D** | mutations of: **W**....**R**.**W**Y **Y D** to A showed defects in taxis toward Ala, Ser and taurine |
| | | | | 5AVE | L-Ser | 3.6 | | |
| | | | | 3C8C | L-Ala | 2.7 | | |
| | | | | 6IOV | L-Arg | 5.6 | | |
| *V. cholerae O395N1* | Mlp24 | WP_001212589.1 | (*36*) | | L_Met | 4.7 | **Y....R.WY Y D** | |
| | | | | 6IOT | L-Arg | 4.8 | | |
| | | | | | L-Ala | 5.1 | | |
| | | | | | L-Gln | 5.5 | | |
| | | | | 6IOR | L-Asn | 6.3 | | |
| | | | | | L-Lys | 6.8 | | |
| | | | | 6IOU | L-Ser | 18.3 | | |
| | | | | | L_Val | 22.1 | | |
| | | | | 6IOS | L-Pro | 23.8 | | |
| | | | | 6IOQ | Gly | 9 | | |
| *P. syringae* pv. actinidiae | PscC | WP_017684350.1 | (*37*) | 6MNI | Pro | 5 ± 0.9 | **Y....R.WY Y D** | |
| *P. syringae* pv. Tomato DC3000 | PscA | WP_011104030.1 | (*38*) | | L-Asp | 1.2 ± 0.1 | **Y....R.WY Y D** | |
| | | | | | L-Glu | 3.4 ± 0.1 | | |
| *P. syringae* pv. Tomato DC3000 | PscA | WP_011104030.1 | (*39*) | | L-Asp | 6.1 | **Y....R.WY Y D** | |
| | | | | | L-Glu | 27 | | |
| *Campylobacter* | Tlp3/CcmL | YP_002344933.1 | (*40*) | 6W3V | L-Phe | 730 ± 55 | **L....K.WY Y D** | mutations of: L....**K**.**W**Y **Y D** to A abolished Ile |

| | | | | | | | | |
|---|---|---|---|---|---|---|---|---|
| *jejuni* subsp. jejuni ATCC 700819 | | | | 4XMR | L-Ile | 86 ± 10 | | binding |
| | | | | | L-Leu | 105 ± 6 | | |
| | | | | | L-Val | 405 ± 27 | | |

**Table S2. Frequencies and variations of the AA_motif found in Bacteria and Archaea in the GTDB representative dataset.** For Bacteria top ten variants are shown. Complete data are in **Table S3**, sheets 3 and 4. NF – not found.

| Motif variant | Protein count in Bacteria | Protein count in Archaea |
|---|---|---|
| Y....R.WY{n1}Y{n2}D | 62.8% | 58.3% |
| F....R.WY{n1}Y{n2}D | 17.2% | 11.3% |
| Y....R.WF{n1}Y{n2}D | 7% | 7.0% |
| W....R.WY{n1}Y{n2}D | 3.7% | NF |
| F....R.WF{n1}Y{n2}D | 2.9% | 23.5% |
| Y....K.WY{n1}Y{n2}D | 1.5% | NF |
| Y....R.WY{n1}F{n2}D | 0.9% | NF |
| F....R.WY{n1}F{n2}D | 0.7% | NF |
| L....R.WY{n1}Y{n2}D | 0.6% | NF |
| W....R.WF{n1}Y{n2}D | 0.5% | NF |

2. Ligand binding studies

We have repeated the isothermal titration calorimetry (ITC) experiments with the mutants of PctA chemoreceptor from *P. aeruginosa* PAO1 using a higher ligand concentration and larger injection volumes (**Fig. S1**).

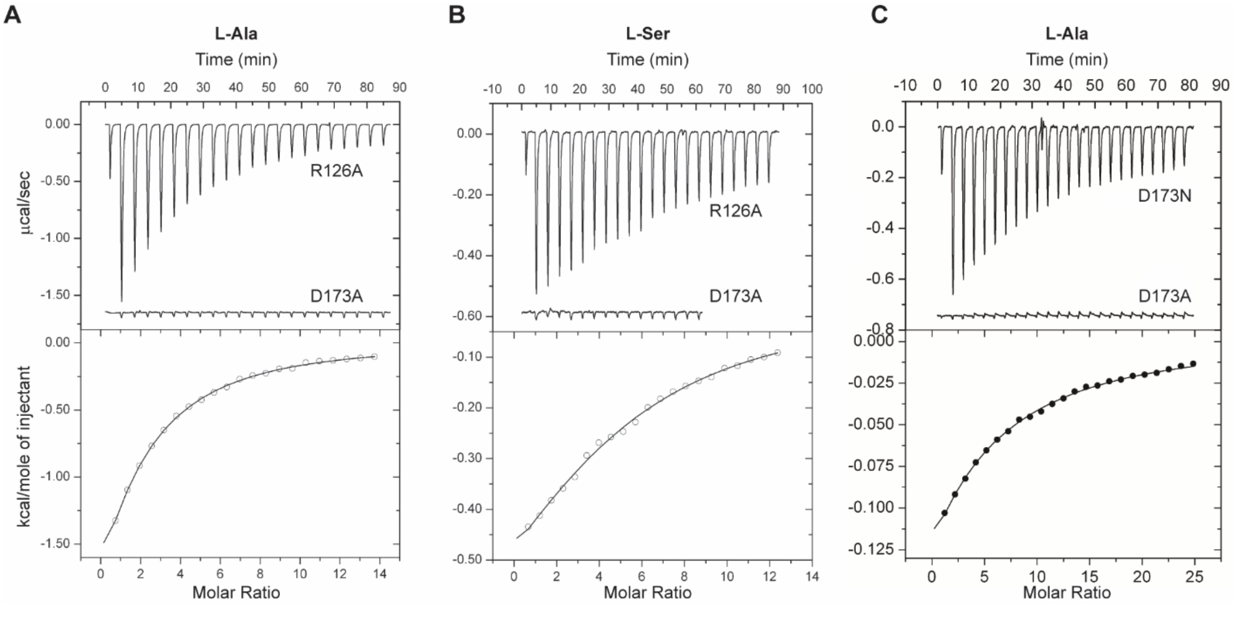


**Fig. S1. Isothermal titration calorimetry binding studies.** (**A**) 3.3 mM L-Ala (12.8 µl injection volume). (**B**) 3 mM L-Ser (12.8 µl injection volume). The protein concentration was 50 µM. R126A and D173A – mutations of two key residues of AA_motif to Ala. (**C**) 15 mM L-Ala (12.8 µl); the protein concentrations were 140 µM and 125 µM for D173N and D173A mutants, respectively. Upper panels: raw titration data. Lower panels: integrated, dilution heat-corrected and concentrationnormalized peak areas of the titration data fitted with the 'One binding site' model of ORIGIN (ORIGINLAB CORPORATION, Northampton, MA, USA).

In the binding study with 3.3 mM L-Ala (12.8 µl injection volume) the D173A mutant has lost its affinity to the ligand (even at higher ligand concentration); the $K_D$ value derived for the R126A mutant was of 202 ± 8 µM, corresponding to a 61-fold reduction as compared to the native protein (**Fig. S1A**). In the study with 3 mM L-Ser (12.8 µl injection volume) similar to L-Ala, D173A was devoid of L-Ser binding and a residual affinity has been determined for the R126A mutant with a $K_D$ value of 258 ± 24 µM (**Fig. S1B**).

In the above experimental set-up, the final ligand concentration in the protein containing sample cell is 175 µM. However, if binding occurred at lower affinity (i.e. $K_D$ values in the mM range) one would not see binding since it does not occur at this low ligand concentration. To drive complex formation and thus to monitor lower affinity binding, the concentration of both ligands must be increased. Therefore, in the next step we conducted the ITC experiments using 30 times more ligand concentration. The upper trace **Fig. S1C** corresponds to a titration of 140 µM PctA D173N mutant with 12.8 µl aliquots of 15 mM L-Ala. Data indicate binding and a $K_D$ constant of approximately 2.1 mM was estimated (important: in **Fig. S1C** for $K_D$ of 1 mM it is better to use the term "estimated" instead of "determined"). We have already shown that the D173A lacks binding activity. To confirm this the PctA D173A mutant was titrated using the same procedure as above (titration of 125 µM protein with 12.8 µl aliquots of 15 mM L-Ala). As shown in the lower trace in **Fig. S1C** no binding heats were observed with high ligand concentration confirming the absence of binding.

Thus, replacement of D173 by A abolishes binding whereas substitution for N reduces the affinity by a factor of approximately 600.

### 3. Identification of the AA_motif in Eukaryota

A search initiated with AA_motif against the Uniprot dataset of dCache_1 domains identified a partial AA_motif in the N-terminal region and the modified AA_motif in the C-terminal region of eukaryotic α2δ proteins. Subsequent MSA of human α2δ proteins with bacterial dCache_1 domains with AA_motif allowed us to discover that the N-terminally located AA_motif is split into two parts by the VWA domain (Pfam Id: PF00092) **(Fig. S2**). Furthermore, the MSA showed that the bacterial dCache_1 domain can be aligned with two regions of human α2δ and CACHD1 proteins **(Fig. S2**): N-terminally and C-terminally located, respectively. The secondary protein structure elements mapped on the alignment showed that the aligned regions of the bacterial and eukaryotic sequences are in good agreement with each other. These data suggested that α2δ and CACHD1 proteins contain two dCache_1 domains and instigated our further investigation.

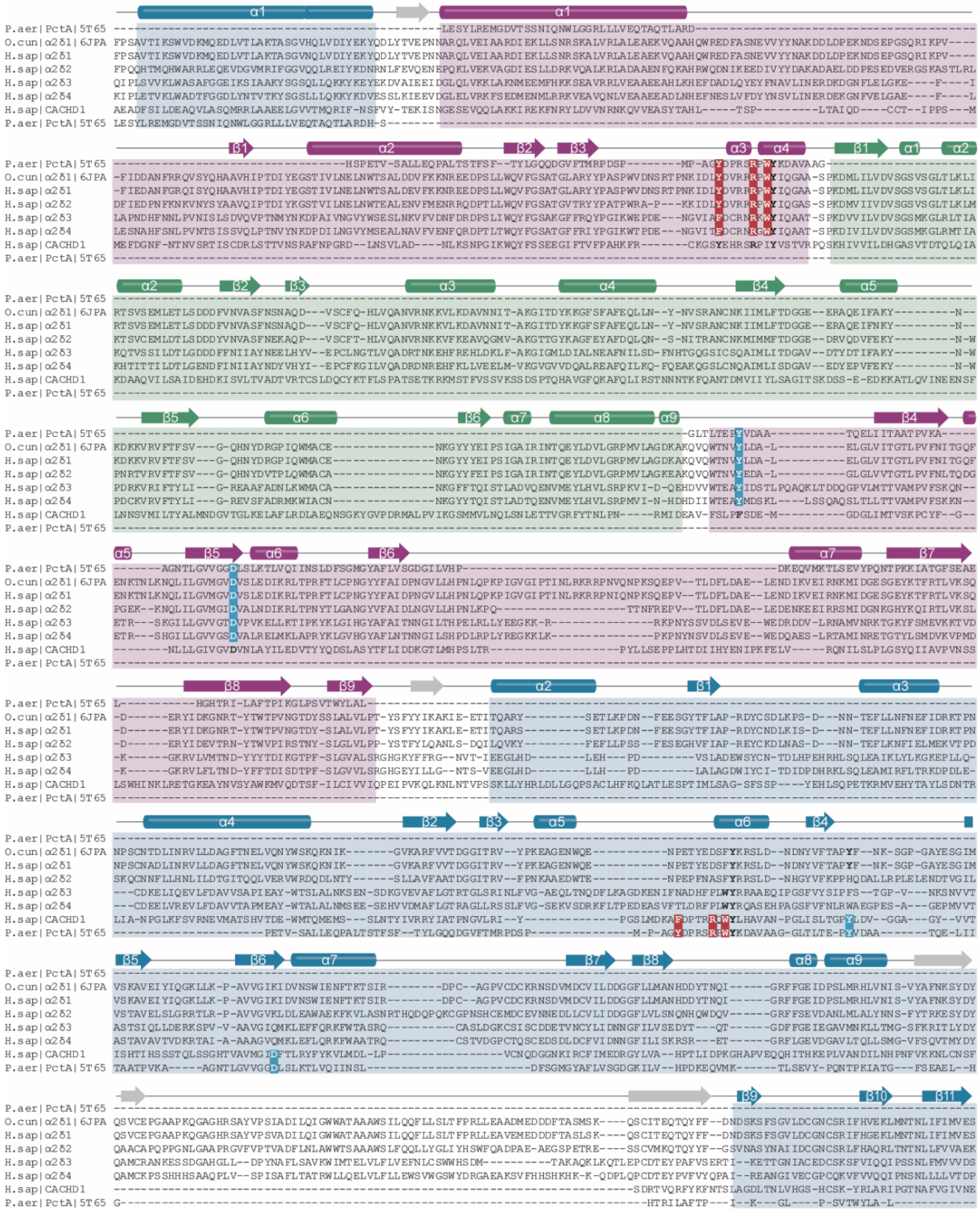

**Fig. S2. Structure-based protein sequence alignment of PctA dCache_1 domain with two regions of the rabbit α2δ-1 and human α2δ and CACHD1 subunits.** Topology is based on the protein structures of PctA (PDB ID 5T65) and α2δ-1 from rabbit (PDB ID 6JPA). 1st dCache_1 domain is shaded in purple, 2nd dCache_1 – in blue, and VWA domain – in green. The AA_motif amino acid positions are in bold and colored in accordance with **Fig. 1**. P.aer – *P. aeruginosa* PAO1, PctA, XP_011514873.1; O.cun – *Oryctolagus cuniculus*, α2δ-1, NP_001075745.1; H.sap – *Homo sapiens*: α2δ-1 – NP_001353796.1, α2δ-2 – NP_006021.2, α2δ-3 – NP_060868.2, α2δ-4 – NP_758952.4, CACHD1 – NP_065976.3.

4. α2δ-1 subunit domain composition

A careful assessment of the available cryo-EM structure of the rabbit α2δ-1 subunit (PDB id of the rabbit VGGC with the α2δ-1 subunit: 6JPA) clearly shows the presence of two dCache_1 domains and one VWA domain (**Fig. 2C**). Both dCache_1 have a long stalk α-helix α1 followed by the upper distal and lower proximal pockets. Usually in bacteria, the dCache_1 domain is enclosed by two transmembrane regions, one preceding the stalk α1 helix and the second following the membrane-proximal pocket. However, the topology tracking of rabbit α2δ-1 and meticulous MSA analysis showed that the stalk α1 helix of the C-terminal dCache_1 (termed here as 2nd dCache_1) is not followed by distal and proximal pockets, but instead by a stalk α1 helix of the N-terminal dCache_1 (termed here as 1st dCache_1) and then by its partial distal pocket. The next structural element is the VWA domain, which is followed by the remaining part of the 1st dCache_1 distal pocket, proximal pocket and then the sequence proceeds to distal and proximal pockets of the 2nd dCache_1 (**Fig. 2D**). This analysis indicates that the 1st dCache_1 is inserted into the loop between α1 helix and the distal pocket of the 2nd dCache_1. It also confirmed that VWA domain is inserted into the 1st dCache_1 distal pocket.

Next, we performed pairwise structural comparison of the dCache_1 domain from the *P. aeruginosa* PAO1 chemoreceptor PctA with both dCache_1 domains of the rabbit α2δ-1 subunit. Remarkably, the PctA dCache_1 domain aligned very well with both dCache_1 domains of the α2δ-1 subunit (**Fig. S3**). The alignment shows essentially identical topologies of the distal and proximal pockets; however, additional secondary structure elements are present in α2δ-1, especially in 2nd dCache_1 (**Fig. S3B**).

Structural analysis also demonstrated that although the AA_motif in the 1st dCache_1 is split into two parts by the VWA domain insertion between α4 and β4, the fold of the distal pocket is intact and amino acid residues that are constitute AA_motif come together and form the interface matching the one in the PctA chemoreceptor (**Fig. S4A)**.

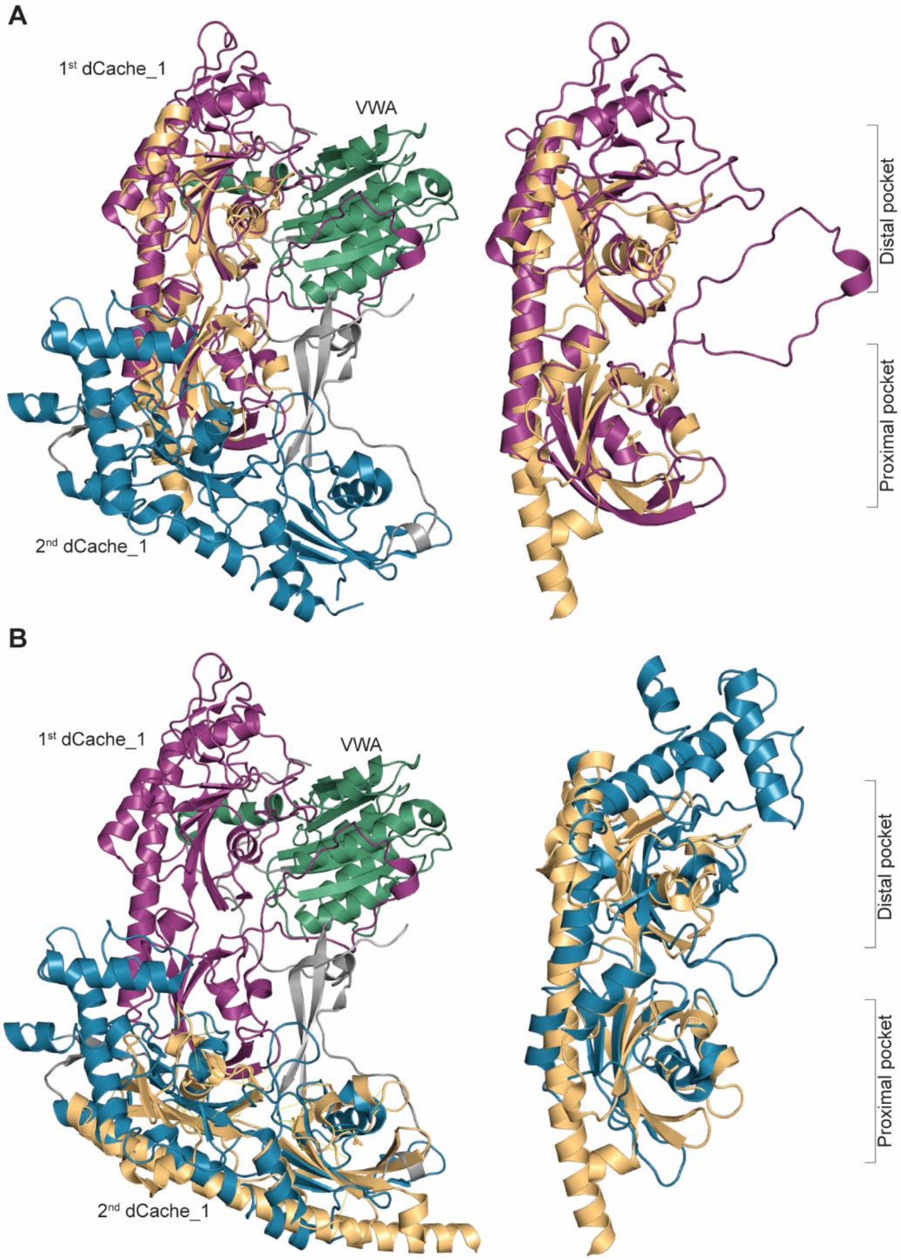


**Fig. S3**. **Superimposition of the *P. aeruginosa* PAO1 PctA dCache_1 (PDB ID 5T65; gold) and two dCache_1 domains of the rabbit α2δ-1 subunit**: (**A**) with the 1st (N-terminally located) dCache_1 domain (purple) and (**B**) with the 2nd (C-terminally located) dCache_1 domain (blue). On the right corresponding superimpositions are enlarged.

5. The ligand-binding pocket of α2δ-1 contains the universal amino acid binding motif

We performed computational docking experiments with the rabbit α2δ-1 subunit and 20 alpha amino acids, GABA, and drug molecules gabapentin, pregabalin, and mirogabalin (see **Materials and Methods**). Using the docking results, we calculated polar contacts made between these ligands and α2δ-1. We established that all these molecules made contacts with the AA_motif residues (Excel **Table S6,** pdb **File 1**). Most ligands made contacts with the first Y, W, the last Y, and the last D of the motif. Less conserved D454 and T463 also made polar contacts with many ligands.

The relative affinities agree with the available data. For example, the docking experiments demonstrated that leucine has higher affinity to α2δ-1 than isoleucine which is consistent with the experiments on inhibition of gabapentin binding by these amino acids (*41*) and experiments measuring specificities of the amino acids (*42*). The order of affinities mirogabalin > gabapentin > pregabalin agrees with recent experimental data (*43*).

6. α2δ-1 subunit and bacterial chemoreceptors bind ligands in a similar fashion

We superimposed the ligand binding pocket (LBP) of the 1st dCache_1 domain of the rabbit α2δ-1 subunit with the LBP of the PctA dCache_1 domain (**Fig. S4A**). The LBPs were extremely similar in shape and size, which is astonishing considering the evolutionary time lapsed from bacteria to mammals and the presence of the VWA insertion. Furthermore, the AA_motif resides in the two structures are located at nearly the same positions.

Next, we closely examined the rabbit α2δ-1 subunit LBP with docked L-Ile in comparison with the LBP of the PctA dCache_1 domain in complex with L-Ile (PDB ID: 5T65) (**Fig. 3**; **Table S9** with the AA_motif coordinates). The position and orientation of L-Ile in α2δ-1 and PctA were nearly identical. Furthermore, ligands made polar contacts with the AA_motif in a similar fashion. The amino group of L-Ile forms hydrogen bonds with the third Y and last D of the AA_motif, whereas the carboxyl group is bound by R and W in both 1st dCache_1 domain of α2δ-1 and dCache_1 domain of PctA. (**Fig. 3**). Computational docking experiments also indicated that L-Leu, gabapentin, pregabalin, and mirogabalin are bound through the AA_motif of the α2δ-1 subunit following the same pattern (**Fig.3, Fig. S4B-D**). The first Y and W of the AA_motif coordinate the ligand carboxyl groups and the third Y (except for the L-Leu ligand) and D interacts with amino groups of the ligands. Carboxyl group of pregabalin is additionally stabilized by a hydrogen bond with R of the AA_motif.

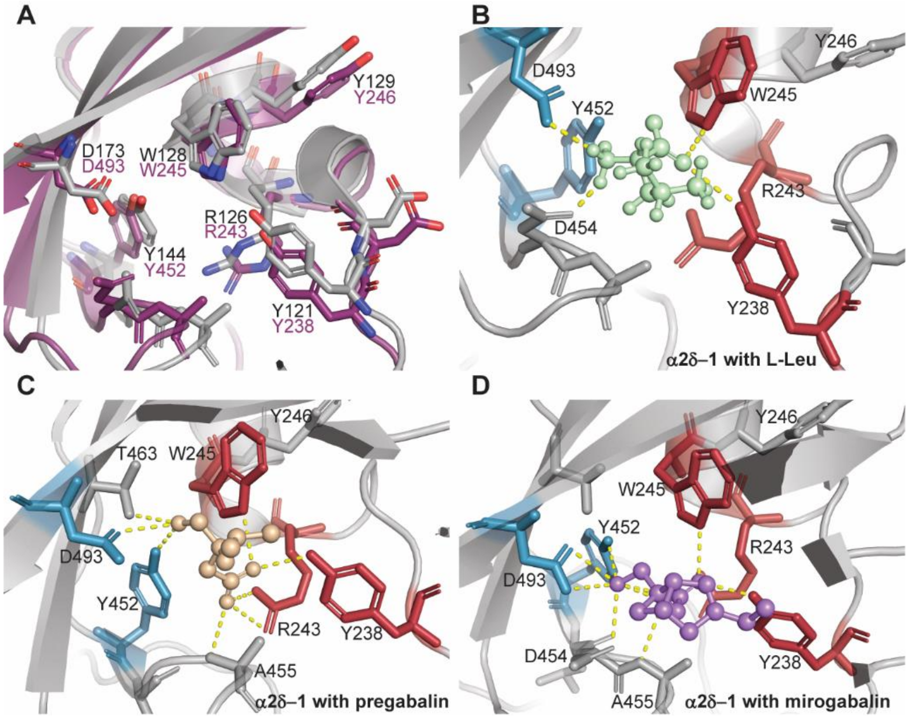


**Fig. S4. Ligand binding pocket (LBP) of the α2δ-1 1st dCache_1 domain.** (**A**) Structural overlay of

the AA_motifs from α2δ-1 1st dCache_1 (purple) with *P. aeruginosa* PctA dCache_1 (grey, PDB ID: 5T65). (**B-D**) LBP of the α2δ-1 1st dCache_1 domain docked with L-Leu (**B,** light green), pregabalin (**C,** beige), and mirogabalin (**D,** violet). The first Y, R and W of the AA_motif are shown in red, and third Y and D are shown in blue.

**Table S9**. **Corresponding amino acid residues of the AA_motif in the rabbit α2δ-1 and in chemoreceptors of *P. aeruginosa* PAO1.**

| **AA_motif residue** | **α2δ-1** (rabbit) | **PctA/PctB** (bacterium) | **PctC** (bacterium) |
|---|---|---|---|
| **Y**….R.WY[n1]Y[n2]D | Y238 | Y121 | Y124 |
| Y….**R**.WY[n1]Y[n2]D | R243 | R126 | R129 |
| Y....R.**W**Y[n1]Y[n2]D | W245 | W128 | W131 |
| Y....R.WY[n1]**Y**[n2]D | Y452 | Y144 | F147 |
| Y....R.WY[n1]Y[n2]**D** | D493 | D173 | D176 |

7. CACHD1 has the same domain architecture as α2δ-1 subunit

Structural modeling using I-TASSER server was performed for the human CACHD1 protein, in which the AA_motif was identified. I-TASSER server uses PDB database to find a template for the best structural match. Structural alignment of the produced CACHD1 model with the rabbit α2δ-1 cryo-EM structure demonstrated that CACHD1 has the same structural composition as α2δ-1 subunit, with a few differences **(Fig. 1, Fig. S5**). Similar to the α2δ-1 subunit, CACHD1 consists of two dCache_1 domains, one inserted into the other one, and VWA domain inserted into the 1st dCache_1 domain. The presence of these structural parts was also confirmed by the MSA (**Fig. S2**). However, α2δ proteins possess distinctive long insertion inside the 2nd dCache_1 between α9 helix and β9 sheet, whereas it is significantly shorter in CACHD1 protein (**Fig. S2**). Another essential difference between α2δ and CACHD1 is the AA_motif placement. Unlike α2δ proteins that all carry the AA_motif in the 1st dCache_1, CACHD1 protein has an intact AA_motif in the distal pocket of the 2nd dCache_1 (**Fig. S2**).

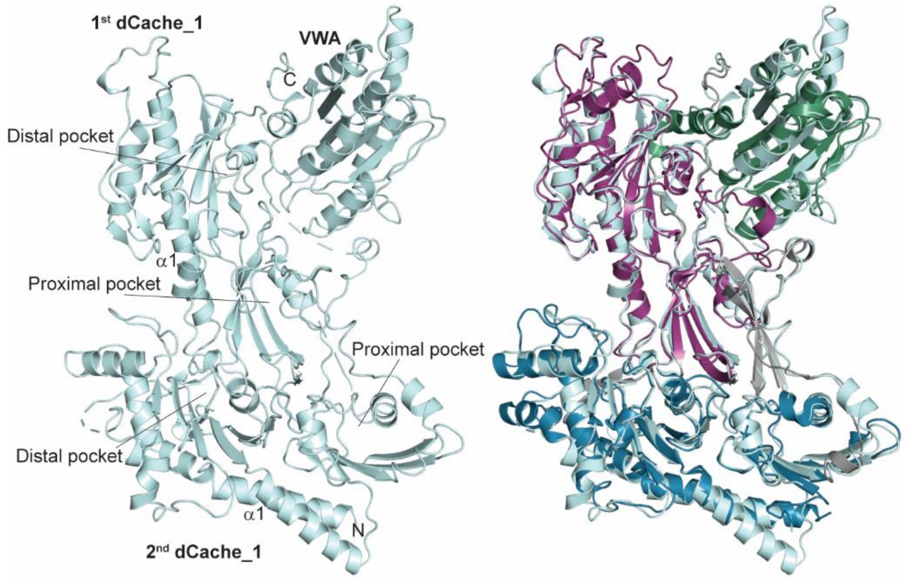


**Fig. S5**. **Structural overlay of the modelled human CACHD1 protein (pale cyan) with the rabbit α2δ-1** (colored as in **Fig. 2**).

8. Evolution of amino acid binding dCache_1 domain in Eukaryota

We found that homologous proteins are present in almost all eukaryotic major groups: Euglenozoa, Heterolobosea, SAR, Haptista, Choanoflagellida, Archaeplastida, and Metazoa (Excel **Table S7,** sheets 1-2**, Fig. 4**). Interestingly, no protein containing dCache_1 domain with AA_motif was found in Angiosperms (flowering plants), whereas they are present in Gymnosperms.

All identified eukaryotic proteins have dCache_1 domain with the inserted VWA domain, which suggests that the VWA insertion probably happened in the Last Eukaryotic Common Ancestor (LECA). Furthermore, our analysis demonstrated that all early diverged branches of eukaryotes (Discoba, SAR, Haptista, Archaeplastida, Cryptophyceae, and Amoebozoa) have proteins with a single dCache_1 domain with the VWA insertion and AA_motif. In contrast, Choanoflagellata and Metazoa have proteins comprised of two dCache_1 domains, one of which has the VWA domain insertion. This indicates that an insertion of the dCache_1-VWA tandem into another dCache_1 domain happened before Choanoflagellata divergence (**Fig. 4**). Of note, choanoflagellate *Salpingoeca rosetta* and some metazoan species have two types of dCache_1 domain containing proteins with AA_motif: one with a single dCache_1 domain with the VWA insertion and one with two dCache_1 domains, one of which has the VWA insertion (**Fig. S7**). Remarkably, some members of Streptophyta clade, for example moss *Physcomitrium patens*, have proteins comprised of one dCache_1 domain with the VWA insertion fused with the serine/threonine kinase domain (Pkinase domain in Pfam nomenclature, PF00069; **Fig. 4**).

We further explored phylogenetic relationships of the proteins with two dCache_1 domains from choanoflagellate and metazoan species. Phylogenetic analysis demonstrated that they fall into three clusters: α2δ, CACHD1, and "double-AA_motif" (**Fig. S6, Fig. S7,** full protein sequence MSA and the tree of eukaryotic double dCache_1 proteins in Newick format can be downloaded from https://github.com/ToshkaDev/Motif repository, "Eukaryotes" folder). α2δ cluster contains only metazoan sequences, including four α2δ proteins from human genome. In Vertebrates α2δ-1 and α2δ-2 proteins form one group, while α2δ-3 and α2δ-4 form another. This suggests that a primordial protein in Vertebrates duplicated, giving rise to two groups of proteins (α2δ-1/α2δ-2 and α2δ-3/α2δ-4 branches) and proteins in each of these groups duplicated one more time. α2δ proteins in bony fish have undergone additional duplications (**Fig. S6, Fig. S7**). *Petromyzon marinus* (sea lamprey) based on the available data does not have proteins in α2δ-1 group but has three proteins in α2δ-2 group.

The CACHD1 cluster includes one protein encoded in the human genome, CACHD1, and proteins from metazoan organisms. (**Fig. S6**). Vertebrates have only one copy of CACHD1, while organisms preceding vertebrates have paralogous proteins in varying numbers (**Fig. S6**, **Fig. S7**).

The third "double-AA_motif" cluster consists of proteins that have AA_motif preserved in the distal pockets of both dCache_1 domains. The proteins in this cluster belong to pre-vertebrate metazoans and Choanoflagellate (**Fig. S7**). Choanoflagellate *S. rosetta* appears to be the first organism to possess double dCache_1 domain proteins (**Fig. 4**), thus indicating that the first double dCache_1 proteins had AA_motif in both dCache_1 domains. Phylogenetic reconstruction suggests that duplications of "double-AA_motif" proteins gave rise to α2δ and CACHD1 clusters (**Fig. S6, Fig. S7**). In α2δ proteins the AA_motif has been preserved predominantly in the 1$^{st}$ dCache_1 domain, while in CACHD1 proteins – in the 2$^{nd}$ dCache_1 domain.

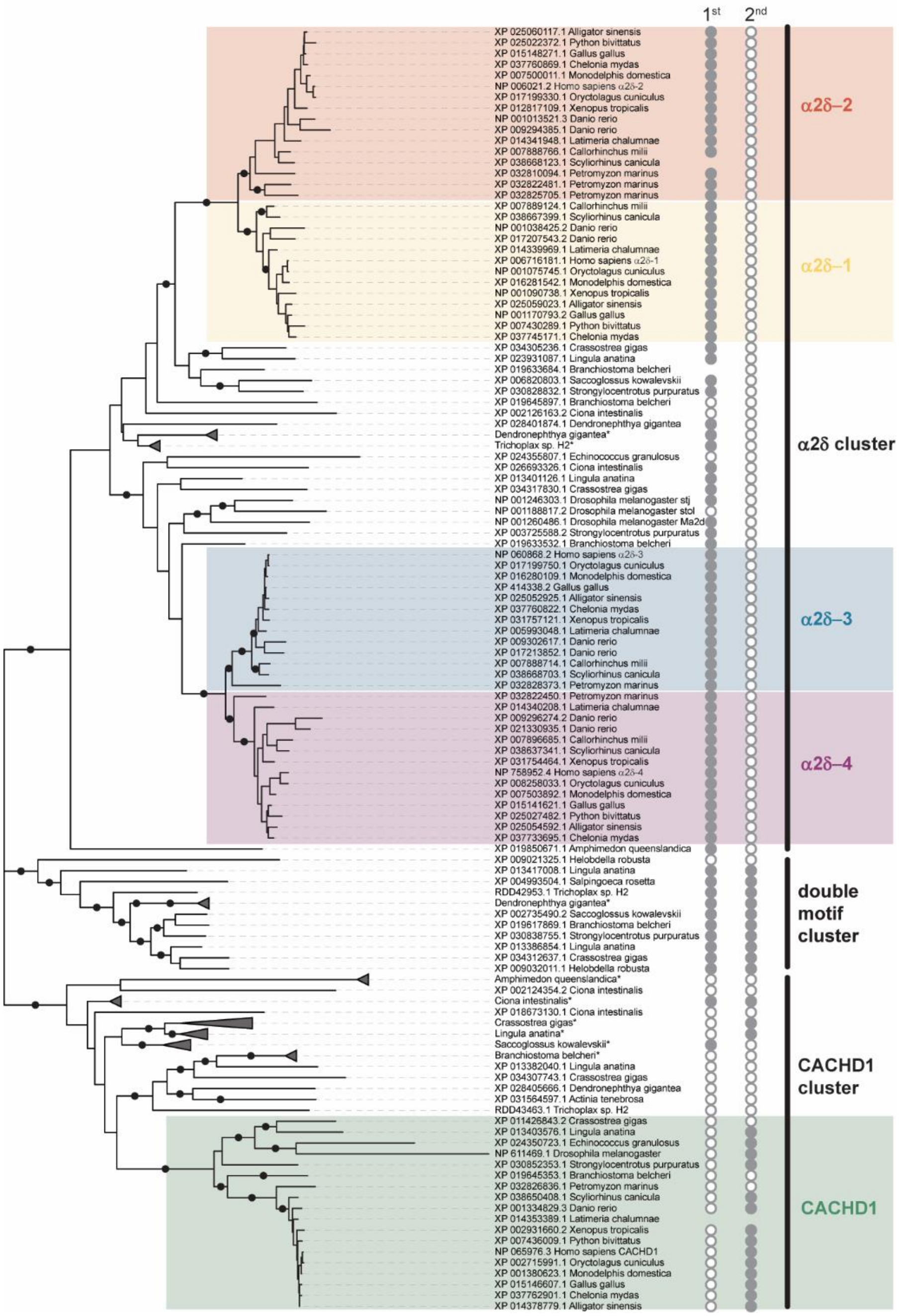


**Fig. S6. Phylogenetic tree of the two dCache_1 domain containing proteins from representative set of eukaryotic species**. Grey circles indicate the presence (filled circle) or the absence (empty circle) of the AA_motif in the 1st and 2nd dCache_1 domains. Black dots mark the bootstrap values that are ≥65. Asterisk after organism names indicates collapsed branches. Full-size figure can be downloaded by this link: https://github.com/ToshkaDev/Motif/raw/main/Eukaryotes/figS6_tree2.tif

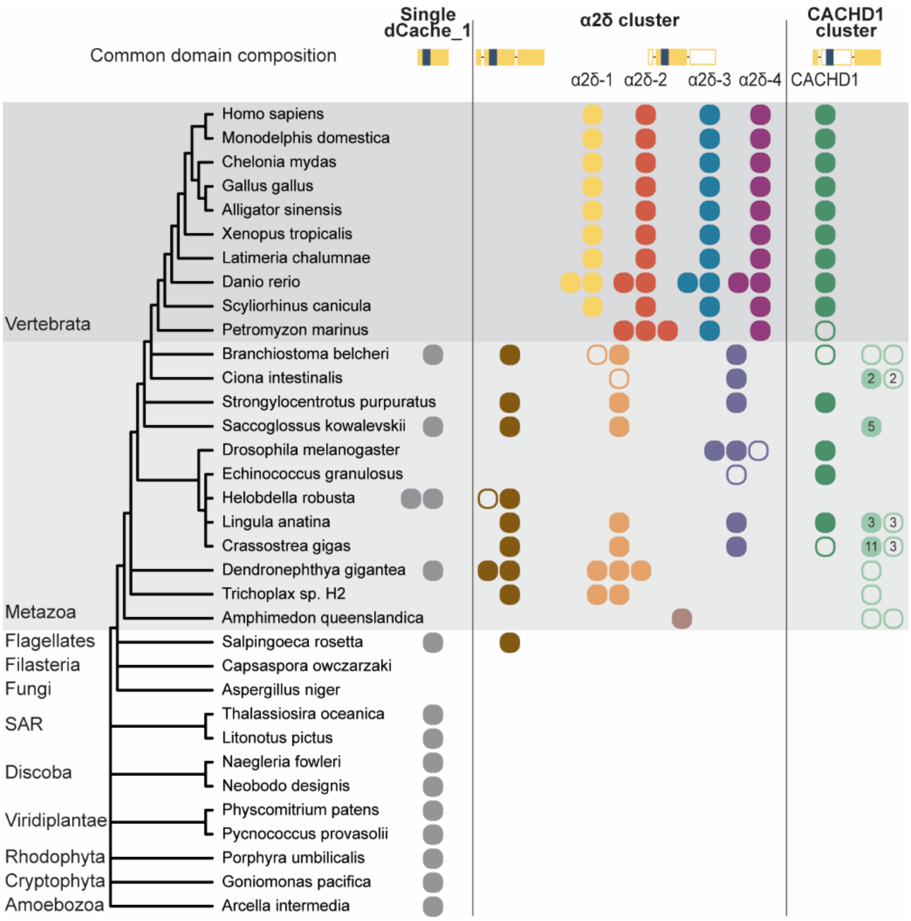


**Fig. S7. Evolutionary history of dCache_1 domain containing proteins across major eukaryotic lineages.** Common domain composition for each family is shown on the top: filled gold rectangles denote dCache_1 domain with the AA_motif, empty gold rectangles – without the motif, dark blue rectangle represents VWA domain. Empty circles denote proteins with no AA_motif in either dCache_1 domain. Numbers inside circles indicate that the genomes of corresponding organisms encode the specified number of paralogous proteins.

<u>9. N-terminal dCache_1 domain of α2δ subunits is under stronger selective pressure than the C-terminal dCache_1 domain</u>

We performed BLASTP and PSI-BLAST searches against prokaryotic proteins in GenBank using several eukaryotic dCache_1 containing proteins with the preserved amino acid binding motif. We initiated separate searches with each of two dCache_1 domains of the proteins. In most cases the top hits were Firmicutes and Proteobacteria (Excel **Table S8**). In almost all searches the coverages of identified proteins were limited to the ligand binding pocket of dCache_1 domains (see Excel **Table S8**). Searches initiated with the 1$^{st}$ dCache_1 domain of α2δ proteins resulted in significant hits. Searches with the human protein easily identified bacterial proteins as significant hits with good E-values. In contrast, the 2$^{nd}$ dCache_1 domain of α2δ proteins from the majority of organisms could not identify significant hits when using BLASTP. Only using PSI-BLAST were we able to identify several significant hits.

Unlike α2δ subunit, both dCache_1 domains of CACHD1 proteins easily found prokaryotic proteins in BLASTP searches. Similarly, searches initiated with both dCache_1 domains of proteins from "double-AA_motif" cluster (**Fig. S6**) readily identified bacterial proteins with significant E-values (see Excel **Table S8**).

19. The PyMOL Molecular Graphics System, Version 2.0, Schrödinger, LLC. (2015).
20. M. Rico-Jimenez *et al.*, Paralogous chemoreceptors mediate chemotaxis towards protein amino acids and the non-protein amino acid gamma-aminobutyrate (GABA). *Molecular microbiology* **88**, 1230-1243 (2013).
21. H. Jeong *et al.*, Genome sequences of Escherichia coli B strains REL606 and BL21(DE3). *Journal of molecular biology* **394**, 644-652 (2009).
22. D. M. Woodcock *et al.*, Quantitative evaluation of Escherichia coli host strains for tolerance to cytosine methylation in plasmid and phage recombinants. *Nucleic Acids Res* **17**, 3469-3478 (1989).
23. J. A. Gavira *et al.*, How Bacterial Chemoreceptors Evolve Novel Ligand Specificities. *mBio* **11**, (2020).
24. K. Taguchi, H. Fukutomi, A. Kuroda, J. Kato, H. Ohtake, Genetic identification of chemotactic transducers for amino acids in Pseudomonas aeruginosa. *Microbiology* **143 ( Pt 10)**, 3223-3229 (1997).
25. J. A. Reyes-Darias, Y. Yang, V. Sourjik, T. Krell, Correlation between signal input and output in PctA and PctB amino acid chemoreceptor of Pseudomonas aeruginosa. *Molecular microbiology* **96**, 513-525 (2015).
26. R. Grist, A. Croker, M. Denne, P. Stallard, Technology Delivered Interventions for Depression and Anxiety in Children and Adolescents: A Systematic Review and Meta-analysis. *Clin Child Fam Psychol Rev*, (2018).
27. J. A. Reyes-Darias *et al.*, Specific gamma-aminobutyrate chemotaxis in pseudomonads with different lifestyle. *Molecular microbiology* **97**, 488-501 (2015).
28. A. Corral-Lugo *et al.*, Assessment of the contribution of chemoreceptor-based signaling to biofilm formation. *Environmental microbiology* **18**, 3355-3372 (2016).
29. B. A. Webb, S. Hildreth, R. F. Helm, B. E. Scharf, Sinorhizobium meliloti chemoreceptor McpU mediates chemotaxis toward host plant exudates through direct proline sensing. *Appl Environ Microbiol* **80**, 3404-3415 (2014).
30. B. A. Webb *et al.*, Sinorhizobium meliloti Chemotaxis to Multiple Amino Acids Is Mediated by the Chemoreceptor McpU. *Mol Plant Microbe Interact* **30**, 770-777 (2017).
31. A. Ud-Din, M. F. Khan, A. Roujeinikova, Broad Specificity of Amino Acid Chemoreceptor CtaA of Pseudomonas fluorescens Is Afforded by Plasticity of Its Amphipathic Ligand-Binding Pocket. *Mol Plant Microbe Interact* **33**, 612-623 (2020).
32. H. Feng *et al.*, Identification of Chemotaxis Compounds in Root Exudates and Their Sensing Chemoreceptors in Plant-Growth-Promoting Rhizobacteria Bacillus amyloliquefaciens SQR9. *Mol Plant Microbe Interact* **31**, 995-1005 (2018).
33. G. D. Glekas *et al.*, A PAS domain binds asparagine in the chemotaxis receptor McpB in Bacillus subtilis. *J Biol Chem* **285**, 1870-1878 (2010).
34. G. D. Glekas *et al.*, Elucidation of the multiple roles of CheD in Bacillus subtilis chemotaxis. *Molecular microbiology* **86**, 743-756 (2012).
35. S. Nishiyama *et al.*, Identification of a Vibrio cholerae chemoreceptor that senses taurine and amino acids as attractants. *Sci Rep* **6**, 20866 (2016).
36. Y. Takahashi, S. I. Nishiyama, K. Sumita, I. Kawagishi, K. Imada, Calcium Ions Modulate Amino Acid Sensing of the Chemoreceptor Mlp24 of Vibrio cholerae. *Journal of bacteriology* **201**, (2019).
37. M. K. G. Ehrhardt, M. L. Gerth, J. M. Johnston, Structure of a double CACHE chemoreceptor ligand-binding domain from Pseudomonas syringae provides insights into the basis of proline recognition. *Biochemical and biophysical research communications* **549**, 194-199 (2021).
38. J. P. Cerna-Vargas *et al.*, Chemoperception of Specific Amino Acids Controls Phytopathogenicity in Pseudomonas syringae pv. tomato. *mBio* **10**, (2019).
